# Pathway profiling of a novel SRC inhibitor, AZD0424, in combination with MEK inhibitors

**DOI:** 10.1101/2021.08.27.457893

**Authors:** John C Dawson, Alison Munro, Kenneth Macleod, Morwenna Muir, Paul Timpson, Robert J Williams, Margaret Frame, Valerie G Brunton, Neil O Carragher

## Abstract

A more comprehensive understanding of how cells respond to drug intervention, the likely immediate signalling responses and how resistance may develop within different microenvironments allows us anticipate how cells adapt to targeted therapy enabling more informed prediction of rational drug combinations. The non-receptor tyrosine kinase SRC regulates many cellular signalling processes and pharmacological inhibition has long been a target of drug discovery projects for the treatment of cancer. Here we describe the *in vitro* and *in vivo* characterisation of the small molecule SRC inhibitor, AZD0424. We show that AZD0424 potently inhibits the phosphorylation of tyrosine-416 of SRC (IC50 ∼ 100 nM) in many cancer cell lines; however inhibition of cell viability, via a G1 cell cycle arrest, was observed only in a sub-set of cancer cell lines in the low (on target) micromolar range. We profiled the changes in intracellular pathway signalling in cancer cells following exposure to AZD0424 and other targeted therapies using Reverse Phase Protein Array analysis. We demonstrate that SRC is activated in response to MEK inhibitor (trametinib or AZD6244)-treatment of KRAS mutant colorectal cell lines (HCT116 and DLD1) and that AZD0424 abrogates this. Cell lines treated with trametinib or AZD6244 in combination with AZD0424 revealed reduction of EGFR, FAK and SRC compensatory activation, and, synergistically inhibits cell viability *in vitro. In vivo*, trametinib-treatment of mice bearing HCT116 tumours increased phosphorylation of SRC on Tyr416, and when combined with AZD0424, inhibition of tumour growth is greater than trametinib alone. We also demonstrate that drug-induced resistance to trametinib is not re-sensitised by AZD0424 treatment *in vitro*, likely as a result of multiple compensatory signalling mechanisms; however inhibition of SRC remains an effective way to block invasion of trametinib resistant tumour cells. These data imply that inhibiting SRC may offer a useful addition to MEK inhibitor combination strategies.

## Introduction

New targeted therapies have been heralded as ‘smart drugs’ that can be tailored to specific cancer subtypes without the adverse toxicity associated with standard chemotherapies. However, clinical studies of many targeted agents in solid tumours have generally failed to produce durable clinical responses, or cure, largely due to compensatory and redundancy mechanisms operating in complex tumours (1). Thus, combinations of targeted agents may be more effective in treating solid tumours, assuming we can identify the signalling networks, often termed rewiring, that permit cancer cells to subvert the activity of single agents. Understanding dynamic compensatory by-pass signalling mechanisms may be able to guide rational drug combinations; with the recent advances in sensitivity, throughput and resolution of transcriptomic and proteomic technologies, we are beginning to understand how chronic drug exposure rewires tumour cell signalling to permit survival. c-SRC (hereafter SRC) is the well known prototype of a large family of non-receptor tyrosine kinases that promotes cancer cell migration, invasion, proliferation, and survival in different contexts (2, 3). SRC activation is widely observed in many types of cancer, such as in solid tumours arising from the colon, breast, lung, liver and pancreas. While SRC is rarely mutated in cancer, it often functions downstream of oncogenic drivers in signalling cascades including those initiated by receptor tyrosine kinases (RTKs) and at integrin-linked focal adhesions (4). SRC has been a target for drug discovery projects for decades with multiple small molecule ATP-competitive inhibitors being tested in clinical trials (2, 5-11). Dasatinib, a multi-kinase SRC inhibitor, is currently approved for the treatment of chronic myeloid and acute lymphoblastic leukaemias (12). While phase I clinical trials have shown that most SRC inhibitors are well tolerated as single agents, trials have generally failed to show significant benefit in advanced solid cancers such as colorectal (for example, (9)), despite strong implicating evidence from preclinical data (13-18). It is therefore becoming evident that preclinical models are failing to predict SRC inhibitor clinical efficacy, most likely because tumour cells are not solely dependent on SRC activity for survival and they can switch to other models of survival and growth signalling. Thus, more unbiased investigations of drugs across genetically distinct cancer cell models, incorporating 2-dimensional(D) and 3-D cell culture and in vivo systems, at both phenotypic and pathway network levels are needed to demonstrate drug sensitivity and resistance, and drug synergies.

Advanced solid tumours, such as metastatic colorectal cancer (CRC) are more likely to harbour combinations of activating mutations in oncogenic driver genes coupled with loss-of-function of tumour suppressor genes (19); therefore, it is not surprising that a single agent targeted therapy is unlikely to succeed. In addition, the tumour microenvironment can influence how tumour cells respond to targeted therapy (20, 21). Such genetic and environmental factors may better be overcome by using combinations of anti-cancer agents that target additional, compensatory or parallel signalling pathways (14, 15, 17, 21).

EGFR is overexpressed in approximately 80% of CRCs and correlates with increased propensity to metastasis and decreased patient survival (19, 22) and EGFR-targeted therapeutic monoclonal antibodies, cetuximab and panitumumab, are approved for the treatment of metastatic disease (23). However, 35-40% of CRC patients have activating mutations of RAS, most frequently of codons 12 or 13 of the KRAS isoform (24). Mutation of KRAS bypasses EGFR signalling, nullifying anti-EGFR targeted therapy, and so patients with CRC tumours harbouring mutant RAS do not generally receive anti-EGFR therapy (25, 26). Cells treated with drugs targeting oncogenic RAS-RAF-MEK signalling can also exhibit inherent and acquired resistance (27). Mechanisms in different contexts include, reactivation of EGFR signalling following MAPK pathway blockade (23, 28, 29), reactivation of MAPK pathway itself (30), activation of parallel pathways (e.g. HER2, MET, or PI3K)(31), or activation of focal adhesion kinase (FAK) to promote tumour cell survival (32). Therefore, understanding how tumour cells respond to putative targeted therapies over time is important to predict how tumour cells escape and survive specific therapies, and guide rational combination hypotheses for clinical testing.

AZD0424 is an orally available potent inhibitor of SRC and ABL with *in vitro* kinase inhibition of ∼4nM (6). In a Phase I clinical trial SRC inhibition was achieved with daily doses ≥ 20mg AZD0424, though no responses were observed as a single agent and only 7 patients (16%) achieved stable disease of 6 weeks or more (6). In this report, we characterise the effect of SRC inhibiton by AZD0424 across preclinical models of breast, prostate and CRC cell lines. We demonstrate that AZD0424 induces a G1 cell cycle arrest in senstitive tumour cell lines, but it does not induce apoptosis. Using Reverse Phase Protein Array (RPPA) analysis, we found that SRC signalling is activated in response to MAPK pathway inhibition by MEK inhibitors in HCT116 CRC cells. We show that in HCT116 cells simultaneous combination of MEK and SRC inhibitors can synergise to reduce cell viability *in vitro* and tumour growth *in vivo*. Finally, we show that while AZD0424 treatment does not resensitise trametinib-resistant HCT116 cells to trametinib treatment with respect to inhibiting cell proliferation, combining AZD0424 and trametinib blocks cancer cell invasion.

## Materials and Methods

### Reagents

All reagents were from Sigma-Aldrich unless otherwise stated. Antibodies were from Cell Signalling Technologies unless otherwise stated.

### Cell Culture

HCT116, HKH2, DLD1 cell lines were provided by S. Van Schaeybroeck (Queens University Belfast, NI). Breast cancer cell lines (BT-549, HCC1954 and SKBR3) were provided by S. Langdon (University of Edinburgh, UK) and MDA-MB-231, PC3, LNCaP, and DU145 cells were purchased from ATCC. Cell lines were cultured in DMEM (Cat No.) supplemented with 10% FCS and 2 mM L-Glutamine. Trametinib resistant HCT116 cells (TRAMR) were generated by treating HCT116 cells with 5 nM trametinib for 2 months in culture and were routinely cultured in 5 nM trametinib thereafter. For experimental drug treatments TRAMR cells were seeded without trametinib.

### Cell viability cell cycle and apoptosis assay

#### Cell seeding and drug treatments

For cell viability assay, 1000-1500 cells were seeded per well of a 96 well plate and grown for 2 days to allow cells to reach exponential growth phase. Media was replaced containing drug with DMSO at a final concentration of 0.1%. Cells were incubated with compounds for 24-72 hours; untreated cells were incubated with 0.1% DMSO.

#### Cell Viability

MTT was added to a final concentration of 1 mg/ml or Alamar Blue added (10-fold dilution) and incubated for 3 hours. For MTT assay, the media was removed, the formazan crystals solubilized in DMSO and, the optical density was measured at 490 nm on a Bio-Rad plate reader. For the Alamar Blue assay and fluorescence emission was read on an EnVision 2101 multilabel plate reader (PerkinElmer; excitation = 540 nm, emission = 590 nm). Results were day 0 subtracted and normalised to control wells for analysis in Prism (Graphpad) to calculate EC50 values using a sigmoidal dose response (variable slope).

#### Cell cycle assay

following drug treatments, cells were fixed in 4 % formaldehyde for 10 minutes, washed three times with PBS and permeabilised with 0.1% Triton X-100 for 5 minutes. Cells were labelled with Hoechst (final concentration 2 μg/mL) for 30 minutes and finally washed three times with PBS. Hoechst labelled nuclei were imaged on a ScanR microscope (Olympus) using a 20× objective, capturing ≥4 fields of view per well. Nuclei were classified into different stages of the cell cycle using the ScanR analysis software.

#### Apoptosis assay

cells were seeded with IncuCyte^®^ Caspase-3/7 green apoptosis assay reagent reagent (Essen BioScience; #4440). Plates were imaged in an IncuCyte Zoom, acquiring images every 3 hours over a 72-hour period using the ‘phase’ and ‘green’ channels. Images were analysed using the IncuCyte Zoom software.

### Organotypic invasion co-culture assay

Organotypic co-cultures were performed as previously described (33). Briefly, dermal fibroblasts were allowed to contract collagen gels over 6-8 days. Tumour cells (4 × 10^4^ cells per gel) were seeded on top of collagen/fibroblast gels. Collagen gels were moved to the air liquid interface on top of a metal grid and allowed to proliferate/invade over a period of 7-9 days. Throughout cells were cultured in 10% FBS/DMEM. Collagen gels were fixed in paraformaldehyde, processed for paraffin embedding and sections cut and stained with H&E.

### Xenograft and in vivo drug treatments

Experiments involving animals were carried out in accordance with the UK Coordinating Committee on Cancer Research guidelines by approved protocol (HO PL 70/8897). For tumour formation, HCC1954 (5 × 10^6^), HCT116 (2 × 10^6^) or DLD1 cells (1 × 10^6^) suspended in Hank’s balanced salt solution, were subcutaneously injected into both flanks of CD-1 Nude mice (Charles River) and allowed to form palpable tumours (>50 mm^3^). Mice were randomised (4/5 per group) and dosed daily by oral gavage with AZD0424, trametinib or a combination made up in 80 mM citrate buffer, pH 3.1, supplemented with 10% Cremaphor EL/10% PEG400. Tumours were monitored twice weekly by calliper measurements and tumour volumes calculated using the following formula *V* = (*W* × *L*)/2, where *V* is tumour volume, *W* is tumour width, and *L* is tumour length. Animals were sacrificed when tumours reached their maximum allowable size or when tumour ulceration occurred. Tumours were fixed overnight in formalin and processed for paraffin embedding and sections cut and stained for H&E using standard techniques.

### Immunohistochemistry

Immunohistochemistry reagents were from DAKO (Agilent Technologies). IHC was performed using standard techniques. Briefly, sections were de-waxed in xylene and antigen retrieval performed in 10 mM citrate buffer using a pressure cooker. Sections were blocked (Peroxidase Block Dako Kit (K4011) and Dako Total Protein Block (X0909)) and incubated with primary antibody overnight (SRC pTyr416 1:200, pERK1/2 1:400). Sections were washed in TBS and incubated with DAB reagent (#K3468) for 5 minutes, and finally, sections were counter-stained with Eosin, dehydrated, and mounted using DPX mounting medium (#44581).

### Western Blotting

Cells were seeded in 6 well plates and allowed to grow over 2 days to ensure cells were in log phase growth. Drug was added in fresh media and cells were lysed in lysis buffer (1% Triton X-100, 50mM HEPES, pH 7.4, 150mM NaCl, 1.5mM MgCl_2_, 1mM EGTA, 100mM NaF, 10mM Na_4_P_2_O_7_, 1mM sodium orthovanadate, 10% glycerol, containing freshly added protease and phosphatase inhibitor cocktails). Lysates were clarified by centrifugation and protein was normalised. Lysates, typically 30µg, was resolved using 4–15% Mini-PROTEAN^®^ TGX™ gels and transferred to Hybond-P 0.45µm PVDF membrane (GE Healthcare). Membranes were blocked in Roche block and incubated with primary antibodies over night or for 3 hours at room temperature. Membranes were washed in TBS-Tween and incubated with anti-rabbit linked HRP secondary antibodies for an hour. Membranes were developed using the BM Chem-Lum substrate (POD) and imaged on a Bio-Rad ChemiDoc MP imaging system. Antibodies were used as per manufacturer’s instructions and listed in Supplementary Table 1.

### Reverse Phase Protein Array (RPPA)

Quantitative Protein expression and phosphorylation profiles were calculated using the Zeptosens reverse phase protein microarray platform as previously described (34). Briefly, cells where rinsed x2 in PBS and lysed in CLB1 buffer (Zeptosens-Bayer) for 30 minutes and centrifuged in microcentrifuge at 14,000 rpm for 5 minutes at room temperature. Supernatants were collected and subjected to total protein determination (coomassie protein assay). Tumour lysates were normalized to a uniform protein concentration with spotting buffer CSBL1 (Zeptosens-Bayer) prior to preparing a final 4-fold concentration series of; 0.2; 0.15; 0.1 and 0.075mg/ml. The diluted concentration series of each sample was printed onto Zeptosens protein microarray chips (ZeptoChip™, Zeptosens-Bayer) under environmentally controlled conditions (constant 50% humidity and 14 °C temperature) using a non-contact printer (Nanoplotter 2.1e, GeSiM). A single 400 Pico litre droplet of each lysate concentration was deposited onto the Zeptosens chip in duplicate spots (thus representing 8 spots per single biological replicate). A reference grid of Alexa Fluor 647 conjugate BSA consisting of 4 columns by 22 rows was spotted onto each sub-array; each sample concentration series were spotted in between reference columns. After array printing, the arrays were blocked with an aerosol of BSA solution using a custom designed nebulizer device (ZeptoFOG™, Zeptosen-Bayer) for 1 hour. Blocked chips were rinsed extensively with water (Milli-Q quality), dried by centrifugation at 200 × g for 5 minutes. Using the built-in micro flow Zeptocarrier system (Zeptosens), the arrays were incubated with different primary antibodies overnight at room temperature. After rinsing the system with assay buffer, the secondary detection antibody (anti-rabbit Alexa Fluor 647) was applied for 2.5 hours at room temperature in the dark. The excess secondary antibody was removed by washing with assay buffer and fluorescence readout of the arrays was performed on the ZeptoReader (Zeptosens) at an extinction wavelength of 635 nm and an emission wavelength of 670 nm. The fluorescence signal was integrated over a period of 1–10 seconds, depending on the signal intensity. Array images were stored as 16-bit TIFF files and analysed with the ZeptoView Pro software package (version 3.1, Zeptosens). Each sample is spotted onto the microarray chip in 2 × 4 dilutions between Alexa Fluor conjugated BSA standards. Fluorescence intensity signals of each sample are calculated by optimized image analysis algorithms and normalized to intensity values of BSA standards through a local 2D quadratic function. A single relative fluorescence intensity (RFI) value is obtained by a weighted linear fit through sample dilutions. RPPA validated antibodies used in the study can be found in Supplementary Table 1.

### Data analysis

For synergy calculations, normalized measurements were averaged (*n* from ≥ 3 independent experiments) and analysed using SynergyFinder (35) using the Bliss, Loewe, and ZIP synergy models.

### Expression analysis

RNA was extracted from HCT116 and TRAMR cells using an RNAse Easy kit (QIAGEN) as per the manufacturers protocol, normalised and equal amounts of the purified RNA, 100⍰ng were used as input for amplification-free RNA quantification by the NanoString nCounter Analysis System with the Human PanCancer pathways panels as previously described(36). Raw counts were normalised to the internal positive controls and housekeeping genes, using the nSolver 4.0 software.

## Results

### *In vitro* characterisation of a novel SRC inhibitor, AZD0424

AZD0424 is an orally available inhibitor of SRC and ABL kinases (*in vitro* SRC kinase IC50 ∼4 nM;(6)), similar to other SRC kinase inhibitors (saracatinib = 2.7 nM, dasatinib = 0.8 nM, bosutinib = 1.2 nM, eCF506 <0.5 nM) (16, 37-39). We sought to first characterise AZD0424 phenotypic and pathway activity across a panel of cancer cell lines with a view to identifying potential indications for drug combination strategies. AZD0424 treatment over 72 hours did not induce potent inhibition of proliferation of the majority of cell lines tested, 11 out of 16 cell lines had EC50 values greater than 5 μM, and of the 5 sensitive cell lines only the colorectal cell line LS174t, displayed an EC50 less than 1 μM (Figure 1A). In our hands LS174t cells displayed high sensitivity to many therapeutic classes in addition to AZD0424, potentially due to reported low expression and high promotor methylation of the ATP⍰binding cassette sub⍰family G member 2 (ABCG2) involved in drug resistance (40). We therefore decided to further characterise AZD0424 in the three sensitive (MDA-MB-231, BT549 and HCC1954) breast cancer cell lines and one that was insensitive (SKBR3). Cell cycle profiling of breast cancer cell lines treated with a range of AZD0424 concentrations for 24 hours revealed, at best, a modest G1-arrest at concentrations greater than 1 μM (Figure 1B) and which is in agreement with other SRC inhibitors such as saracatinib or dasatinib (37, 41). Finally, we observed no change in the induction of apoptosis using an activated caspase 3/7 assay (Figure 1C).

**Figure 1.**
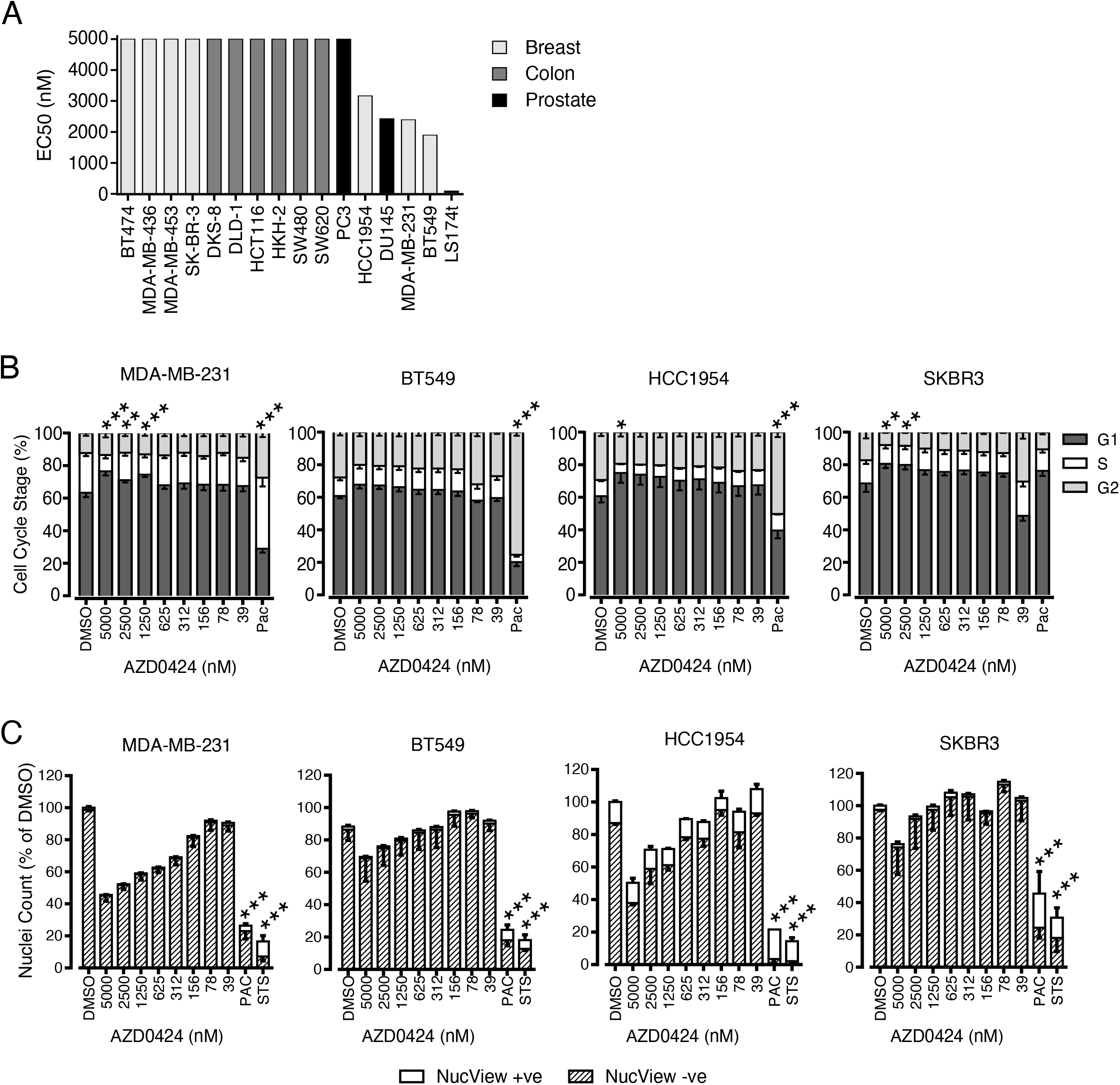
Profiling AZD0424 across cancer cell lines. (A) Ranked AZD0424 EC50 values for cell viability of cancer cell lines treated with AZD0424 (*n* = 3 independent experiments). (B) Cell cycle distribution after 24 hours of AZD0424 treatment. Bars represent mean percentage of cells in each stage of the cell cycle ± standard deviation (*n* = 3 independent experiments). (C) Measurement of nuclei number and apoptosis following 48 hours treatment of breast cancer cell lines with AZD0424. Data are mean ± SEM (*n* = 3 independent experiments). *, *p* < 0.05; **, *p* < 0.01; ***, *p* < 0.001 (two-way ANOVA).

We next determined the ability of AZD0424 to inhibit cellular SRC kinase activity by performing RPPA analysis across compound dose-response and time-series studies performed in the breast cancer cell line panel (Figure 2A, top panel and Supplementary Figure 1). Increasing concentrations of AZD0424 rapidly elevated SRC protein levels within 3 hours of treatment which was sustained over a 24-hour period (Figure 2A). Concomitantly, we observed a concentration dependent decrease in SRC-family kinase activation, as measured by the phosphorylation of Tyr416 (Figure 2A, middle panel and Supplementary Figure 1B). AZD0424 induced SRC inhibition with a cellular EC50 of ∼100 nM. RPPA profiling of AZD0424 response revealed a number of pathway markers that were also inhibited in a concentration dependent manner demonstrating the wider impact of SRC inhibition on cellular signalling (Supplementary Figure 1C). AZD0424 treatment induced reduction of phosphorylation of the SRC kinase target STAT5 (Tyr694) in addition to EGFR family (Tyr1248/Tyr1173), PLCγ (Tyr783), and SHP2 (Tyr542) signalling.

**Figure 2.**
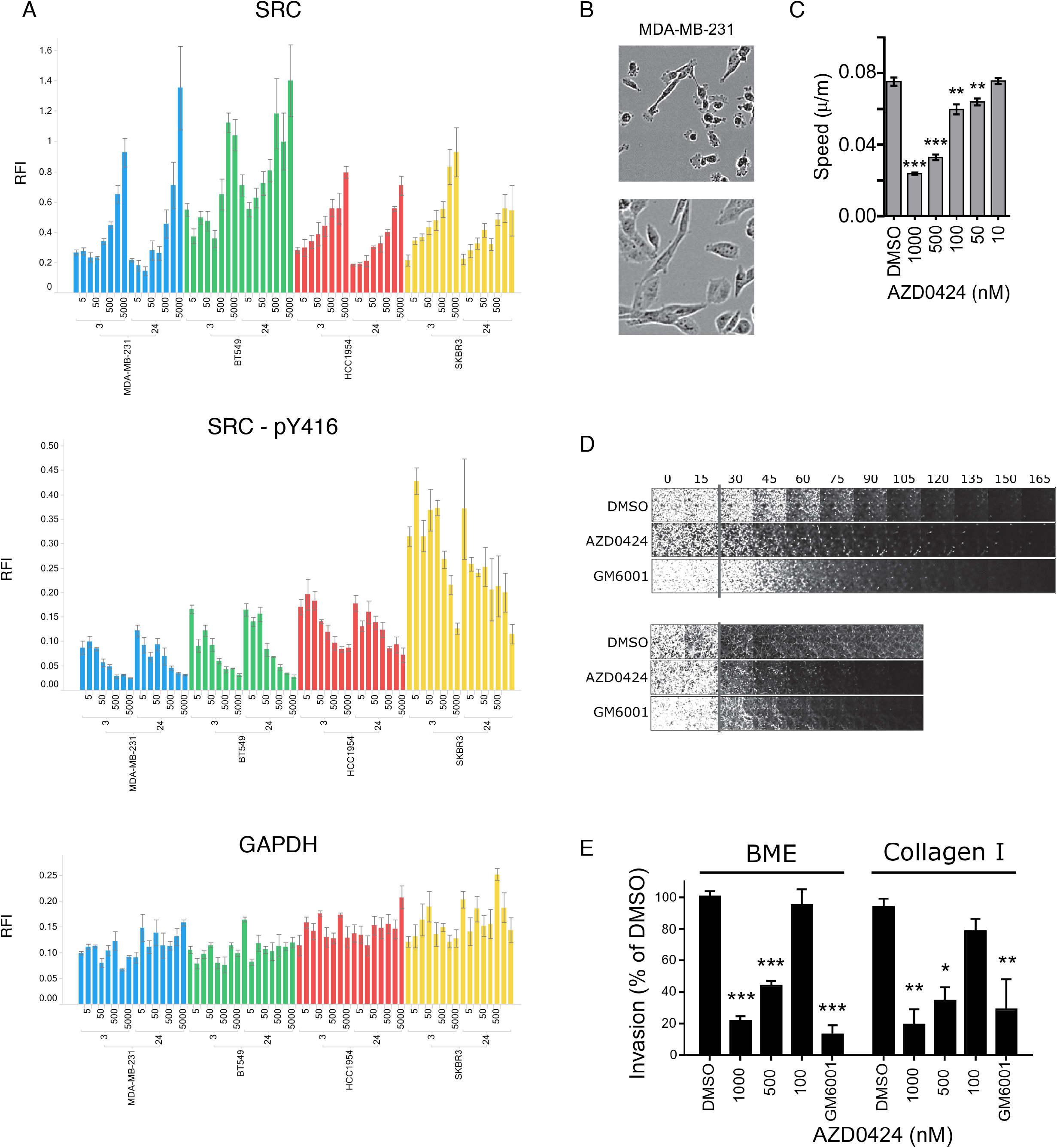
AZD0424 treatment of breast cancer cells. (A) RPPA profiling of total and phosphorylated SRC (Tyr416) in breast cancer cells treated with AZD0424. Mean relative fluorescence intensity (RFI) is shown ± SEM (*n* = 3 technical replicates). (B) AZD0424 inhibits random migration of MDA-MB-231 cells. Graph is mean ± SEM from 3 independent experiments. (C) AZD0424 inhibits the invasion of MDA-MB-231 cells into basement membrane extract (top panels) or collagen I matrix (bottom panels). (D) Quantification of MDA-MB-231 invasion. Data are mean ± SEM (*n* = 3 independent experiments).

SRC is a well-known regulator of cancer cell motility and invasion and so, using the invasive MDA-MB-231 cell line, we tested the ability of AZD0424 to inhibit random migration and invasion into distinct 3D extracellular matrices using basement membrane extract (BME) or collagen I (Figure 2B-D). In 2D cell culture, AZD0424 treatment of MDA-MB-231 cells induced a morphological change with cells becoming flatter with fewer protrusions compared to the control cells (Figure 2B) and reduced the speed of random migration at concentrations (100-500 nM) comparable to on-target cellular SRC inhibition. Further, AZD0424 inhibited the ability of MDA-MB-231 cells to invade through either 3D BME or collagen I matrix (Figure 2D).

Additionally, we tested the ability of AZD0424 to inhibit SRC activation and tumour growth of one of the sensitive breast cancer cell lines (HCC1954) *in vivo*. HCC1954 cells were injected subcutaneously into the flanks of CD-1 Nude mice and tumour bearing mice were dosed with AZD0424 daily (Supplementary Figure 2). AZD0424 did not affect the growth of HCC1954 tumour xenografts even though SRC was effectively inhibited using daily dosing of mice with concentrations of ≥ 10 mg/kg (Supplementary Figure 2B). These studies clearly demonstrate that despite potent inhibition of intracellular SRC activity, AZD0424 has minimal impact upon cancer cell survival in these models.

### AZD0424 as a potential combination therapy in KRAS colorectal cancer

As SRC inhibitors perform poorly as single anti-cancer agents in most cancers tested (for example (9)), we next sought to identify potential resistance mechanisms that rely upon SRC that could be targeted with drug combination therapy using AZD0424. To inform a Phase I clinical trial containing predominantly patients with colorectal cancer (19 patients; 47% of total(6)) for potential combination treatments with AZD0424, we applied RPPA to profile the response of four colorectal cancer cell lines with mutations in the KRAS gene, a common mutation in CRC, to treatment with the MEK inhibitors trametinib and AZD6244. Treatment of HCT116, and to a lesser extent of DLD1 cells, with trametinib or AZD6244 induced the activation of SRC, as measured by an increase in phosphorylation of Tyr416 (Figure 3A-C). In addition, MEK inhibitor treatment also resulted in a compensatory induction in phosphorylation of a number of other proteins involved in EGFR/RTK signalling (Figure 3B and Supplementary Figure 1) including STAT5 (Tyr694), EGFR (Tyr1068/Tyr1173), PLCγ (Tyr783), IGF-1R beta (Tyr1162/Tyr1163) and SHP2 (Tyr542). Notably, the activation of EGFR was greater in the DLD1 cells compared to the HCT116 cells. We next tested whether co-treatment with AZD0424 could inhibit the activation of compensatory signalling induced by either trametinib or AZD6244 in HCT116 or DLD1 cells (Figure 3B). Treatment with either MEK inhibitor reduced the activation of ERK1/2 and also elevated phosphorylation of MEK1 (Ser21/Ser221) itself, and FAK (Tyr397). MEK inhibitor treatment also blocked the activation of S6 ribosomal protein and the cell cycle regulator Rb, characteristic of a cell cycle arrest. HCT116 and DLD1 cells treated with a combination of AZD0424 and a MEK inhibitor blocked the activation of EGFR, SHP2, PLCγ, but not FAK (Tyr397) suggesting that reactivation of signalling through the EGFR pathway in response to MEK inhibition requires SRC activity while the activation of FAK by autophosphorylation on Tyr397 is independent of SRC. MEK inhibitor treatment of HCT116, and to a lesser extent DLD1 cells, increased the phosphorylation of FAK on Tyr861, a SRC kinase substrate, which could be inhibited by AZD0424 treatment (Figure 3C and D). Interestingly, the pattern of compensatory pathway signalling in response to MEK inhibitor treatment is cell type dependent and HCT116 and DLD1 cells displayed differences in their response to MEK inhibitor treatment, for example HCT116 cells did not activate the EGFR pathway or activate AKT (phosphorylation of Ser473) as strongly as DLD1 cells (Figure 3B and 3C).

**Figure 3.**
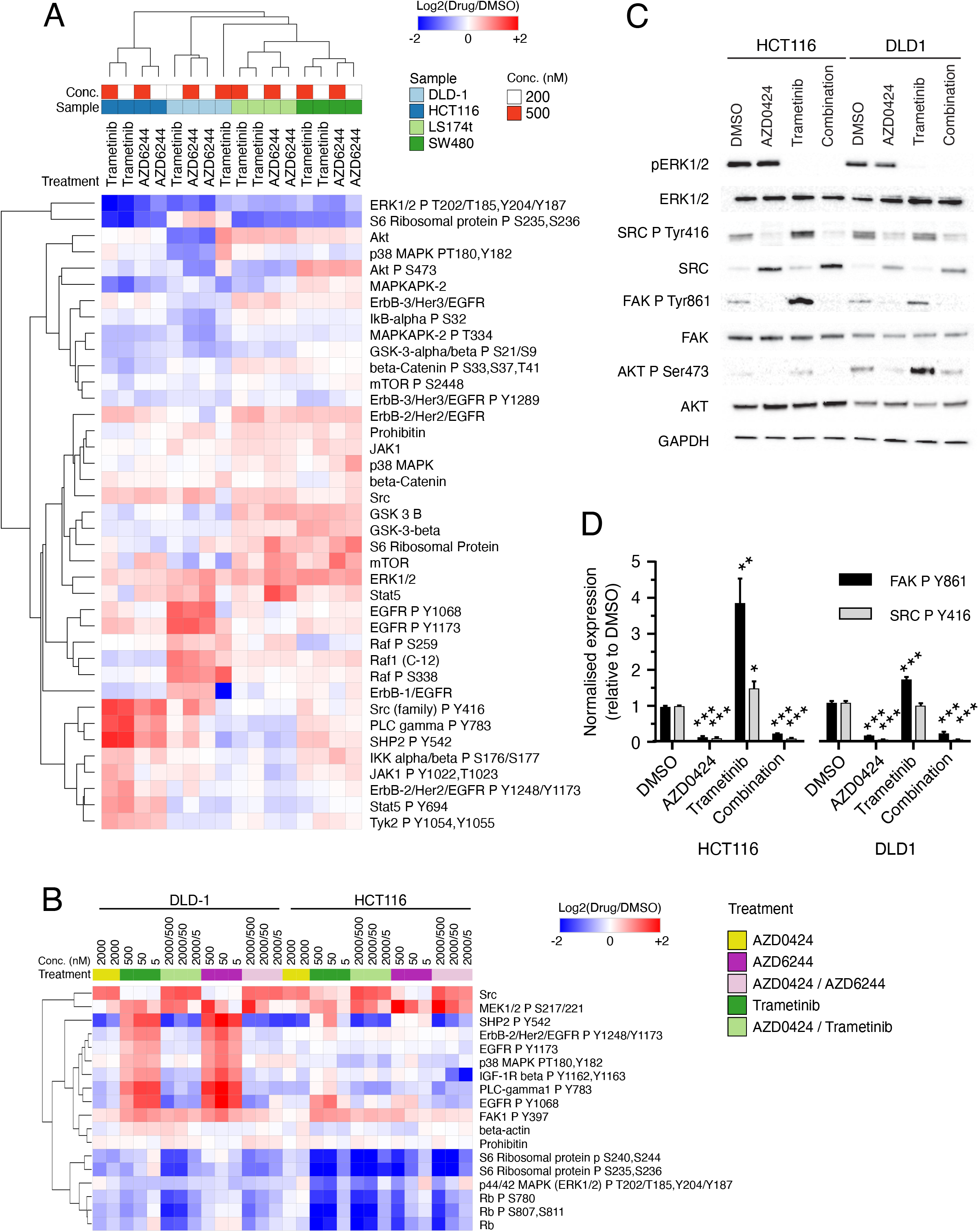
MEK inhibitors activate SRC signalling in KRAS-G13D mutant colorectal cancer cell lines. (A) Heatmap showing RPPA profiling of colorectal cell lines treated with MEK inhibitors for 24 hours. (B) RPPA profiling of HCT116 and DLD1 cell lines treated with AZD0424 alone or in combination with the MEK inhibitors trametinib or AZD6244 for 24 hours. In both (A) and (B), hierarchal clustering using Euclidean distance and complete linkage is shown. Values are normalised to DMSO treated samples. (C) Western blot analysis of signalling changes following treatment with AZD0424 (2000 nM) and or trametinib (5 nM) treatment for 24 hours. (D) Quantification of western blot changes for phosphorylated SRC Tyr416 and FAK Tyr861. Mean ± SEM is shown (*n* = 3 independent experiments). *, *p* < 0.05; **, *p* < 0.01; ***, *p* < 0.001 (one-way ANOVA).

### AZD0424 synergises with MEK inhibitors in two KRAS-G13D mutant colorectal cancer cell lines

Having confirmed that AZD0424 might block potential compensatory signalling induced by MEK inhibitors in KRAS mutant HCT116 and DLD1 cells, we next tested whether the combination could reduce cell viability (Figure 4A). Measurement of cell viability following inhibitor treatment revealed that DLD1 cells were resistant to MEK inhibitor treatment (trametinib, EC50 >300 nM; AZD6244, EC50 > 3000 μM), HCT116 cells in contrast were sensitive (trametinib, EC50 = 1.5 nM; AZD6244, EC50 = 127 nM) consistent with previous reports (30, 31). Conversely, DLD1 cells were more sensitive to AZD0424 treatment than HCT116 (Figure 4A), though at much higher concentrations (3μM) than that required to inhibit cellular SRC (Figure 2A). Treatment of cells with AZD0424 in combination with either trametinib or AZD6244 resulted in synergistic inhibition of cell viability at sub-μM doses in both cell lines (Figure 4A and B). Finally, we confirmed that SRC inhibition using dasatinib in combination with either trametinib or AZD6244, also resulted in a synergistic inhibition of cell viability in both cell lines (Supplementary Figure 3A-C).

**Figure 4.**
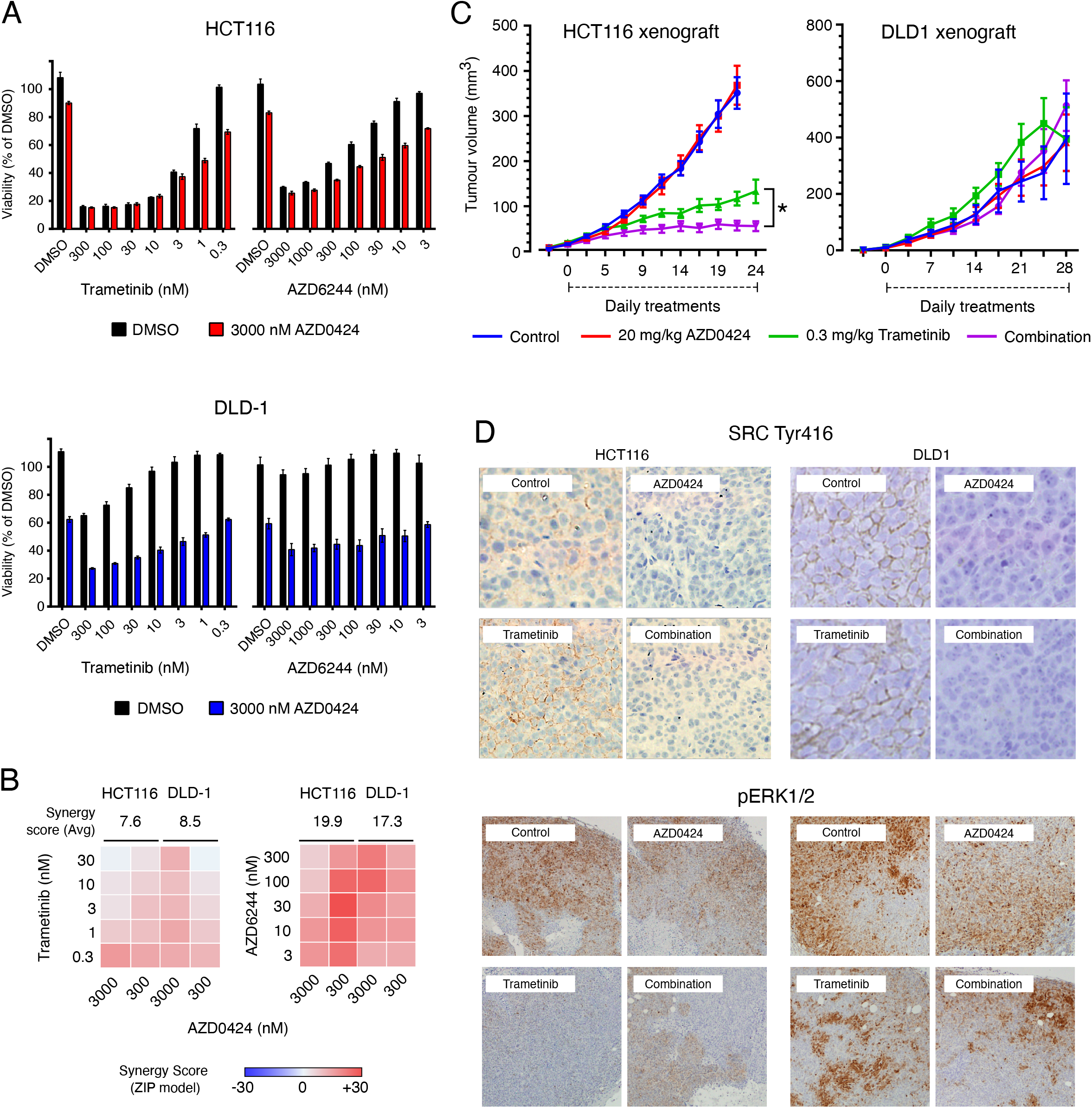
MEK and SRC inhibitors synergistically inhibit proliferation. (A) AZD0424 synergistically inhibits cell viability of DLD1 and HCT116 cells in combination with trametinib or AZD6244. Mean cell viability is shown ± SEM (*n* = 3 independent experiments). (B) Synergy analysis for trametinib or AZD6244 in combination with AZD0424. Synergy was calculated using the ZIP synergy model. (C) Combination of AZD0424 and trametinib inhibits HCT116 (left) but not DLD1 (right) tumour growth. Tumour volumes are plotted as means ± SEM [*n* ≥ 4 mice per group (2 tumours per mouse)]. *, p < 0.05 (one-way ANOVA). (D) Immunohistochemical analysis of phosphorylated SRC Tyr416 and ERK1/2 from 5-day drug treated tumours.

We next tested whether AZD0424 combined with MEK inhibitors could inhibit tumour growth in xenograft models using both DLD1 and HCT116 cells. Mice bearing tumours formed by injecting HCT116 cells subcutaneously on the flanks of CD-1 Nude mice were dosed by oral gavage daily with trametinib (Supplementary Figure 3D). Trametinib was very effective at blocking HCT116 tumour growth in a concentration dependent manner and so we selected 0.3 mg/kg as a dose of trametinib that did not achieve complete growth inhibition to test in combination with AZD0424 (Figure 4C). Treatment with AZD0424 alone had no effect on HCT116 or DLD1 tumour growth, while HCT116 tumours treated with AZD0424 in combination with 0.3 mg/kg trametinib showed a significant reduction of tumour growth compared to trametinib alone (Figure 4C). Strikingly, DLD1 tumours were not only resistant to treatment with AZD0424, but also to trametinib alone and in combination (Figure 4C), despite strong *in vitro* synergy (Figure 4A and B). We confirmed that trametinib treatment activates SRC in HCT116 tumours *in vivo* and that SRC activation is effectively inhibited using AZD0424 alone or in combination with trametinib (Figure 4D). Trametinib, at 0.3 mg/kg, only partially inhibited ERK1/2 activation as expected at this submaximal dose in both HCT116 and DLD1 tumours and the addition of AZD0424 did not alter this (Figure 4D). DLD1 tumours treated with trametinib also exhibited pockets of strong pERK1/2 staining which appeared undiminished by trametinib, alone or in combination with AZD0424. As DLD1 cells are resistant to MEK inhibitors and display a stronger activation of EGFR signalling upon treatment with MEK inhibitors, we hypothesised that DLD1 cells were more dependent on EGFR signalling for survival. We therefore tested DLD1 cells with drug combinations targeting MEK inhibition (trametinib) and either EGFR (AZD8931 and Lapatinib) signalling or AKT (AZD5363), as this cell line also has an activating mutation in the *PI3KCA* gene (amino acid E545K) (Supplementary Figure 3E and F). Combined treatment of trametinib with AZD5363, AZD8931, or lapatinib synergistically inhibited DLD1 cell viability. Interestingly, this could be further enhanced by the addition of AZD0424 as a triple combination suggesting the involvement of SRC signalling (Supplementary Figure 3E and F).

### AZD0424 does not sensitise MEK inhibitor-resistant cells to MEK inhibition

As AZD0424 did not sensitise DLD1 tumours that were inherently resistant to MEK inhibitor treatment, we next asked whether SRC inhibition could (re)sensitise cells that had acquired resistance to MEK inhibitors following prolonged treatment with trametinib. We generated HCT116 trametinib-resistant cells (TRAMR) by long term exposure to 5 nM trametinib in cell culture and compared their sensitivity to trametinib to the parental HCT116 cells and an isogenic cell line (HKH2), where the copy of the mutant KRAS gene has been deleted (Figure 5A). Both the HKH2 and TRAMR cell lines were less sensitive to trametinib treatment (EC50 = 5.9 and 28 nM, respectively) compared with parental HCT116 cells (1.2nM) as measured by cell viability (Figure 5A); however, co-treatment with the combination of AZD0424 and trametinib still synergistically inhibited cell viability in the HKH2 and TRAMR cells (Figure 5B). Short term exposure to trametinib (24 hours) resulted in strong activation of SRC signalling across HCT116, HKH2 and TRAMR cells, as demonstrated by elevated phosphorylation of Src Tyr416 and FAK Tyr861 and this was prevented by co-treatment with AZD0424 (Figure 5C). AZD0424 treatment alone blocked phosphorylation of SRC Tyr416, FAK Tyr861, in all cells and partially ERK1/2 in HKH2 cells (Figure 5C). The basal activation of ERK1/2 was elevated in TRAMR cells and was insensitive to treatment with AZD0424 and only partially sensitive to trametinib. Drug induced resistance to MEK inhibitors *in vitro* can be driven by amplification of the KRAS gene resulting in re-activation of the MAPK pathway (30, 42), and transcriptomic analysis of the TRAMR cells demonstrated elevated expression of KRAS mRNA and the upregulation of genes in the MAPK, PI3K, JAK-STAT and Wnt pathways (Supplementary Figure 4). To further profile trametinib-induced resistance at the post-translational pathway level, we treated HCT116 and TRAMR cells for 24 hours with AZD0424, trametinib, and AZD6244 and profiled some post translational modifications using RPPA analysis (Figure 5D). Compared to parental HCT116 cells, TRAMR cells had elevated levels of phosphorylated MEK1 (Ser217/Ser221) and ERK1/2 confirming stimulation of the MAPK pathway as a likely mechanism of resistance to trametinib. TRAMR cells also had elevated IRS-1 expression, phosphorylation of AKT (Ser473), and to a lesser extent phosphorylation of GSK3α/β (Ser9, Ser21), p90 S6 kinase (Thr359, Ser363), and Rb (Ser780). Treatment of TRAMR cells with trametinib or AZD6244 increased phosphorylation of EGFR (Tyr1068/Tyr1173), FAK (Tyr397), PLCγ (Tyr783), SHP2 (Tyr542), and STAT5 (Tyr694), as previously observed in HCT116 cells (Figure 3). Further, in TRAMR cells, AZD0424 could effectively block the activation of many of these compensatory signalling proteins but not amplified phosphorylation of; MEK1/2 (Ser217/221), c-Jun (Ser7) and Akt (Ser473) (Figure 5D). Therefore, MEK inhibitor resistance driven by increased MAPK signalling pathway is unlikely to benefit from SRC inhibitor combination therapy alone.

**Figure 5.**
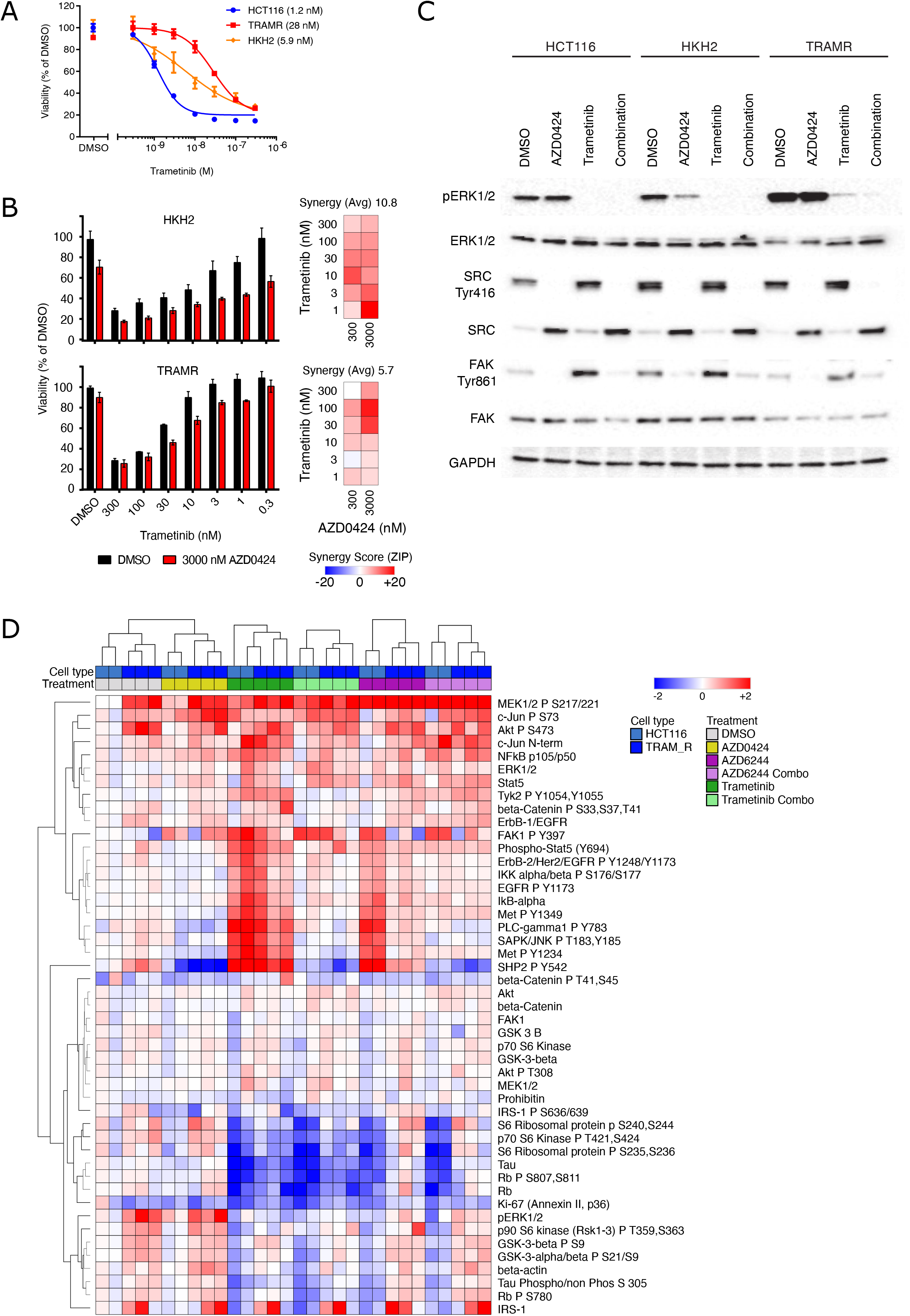
SRC and MEK inhibitor combinations do not sensitise drug-induced resistant cells. (A) Cell viability of cells treated with trametinib. Trametinib resistant HCT116 cells (TRAMR). Isogenic wild-type KRAS HCT116 cells (HKH2). EC50 values shown in parenthesis. (B) Cell viability of HKH2 and TRAMR cells in combination with AZD0424 after 3 days treatment. Mean cell viability is shown ± SEM (*n* = 3 independent experiments). (C) Western blot analysis of lysates from cells treated with AZD0424 (2000 nM) or trametinib (5 nM) for 24 hours. (D) RPPA analysis of lysates from cells treated with AZD0424 (2000 nM), AZD6244 (2000 nM) and trametinib (5 nM) for 24 hours. Hierarchal clustering using Euclidean distance and complete linkage is shown. Values are normalised to DMSO treated samples.

### AZD0424 and trametinib synergistically inhibit cancer cell invasion

SRC and ERK1/2 regulate cancer cell migration, invasion and metastasis (13) and, therefore, we tested whether combinations of MEK and SRC inhibitors could synergise to block invasion using a 3D organotypic collagen I invasion assay (33). As observed in the cell viability assay (Figure 4A), trametinib treatment inhibited proliferation of HCT116, but not DLD1 cells, where combined treatment with trametinib and AZD0424 was required (Figure 6A and B). Both HCT116 and DLD1 cells invaded into organotypic collagen I matrices, and invasion was readily inhibited by treatment with either AZD0424 or trametinib and this was enhanced by combining the two agents (Figure 6A and B).

**Figure 6.**
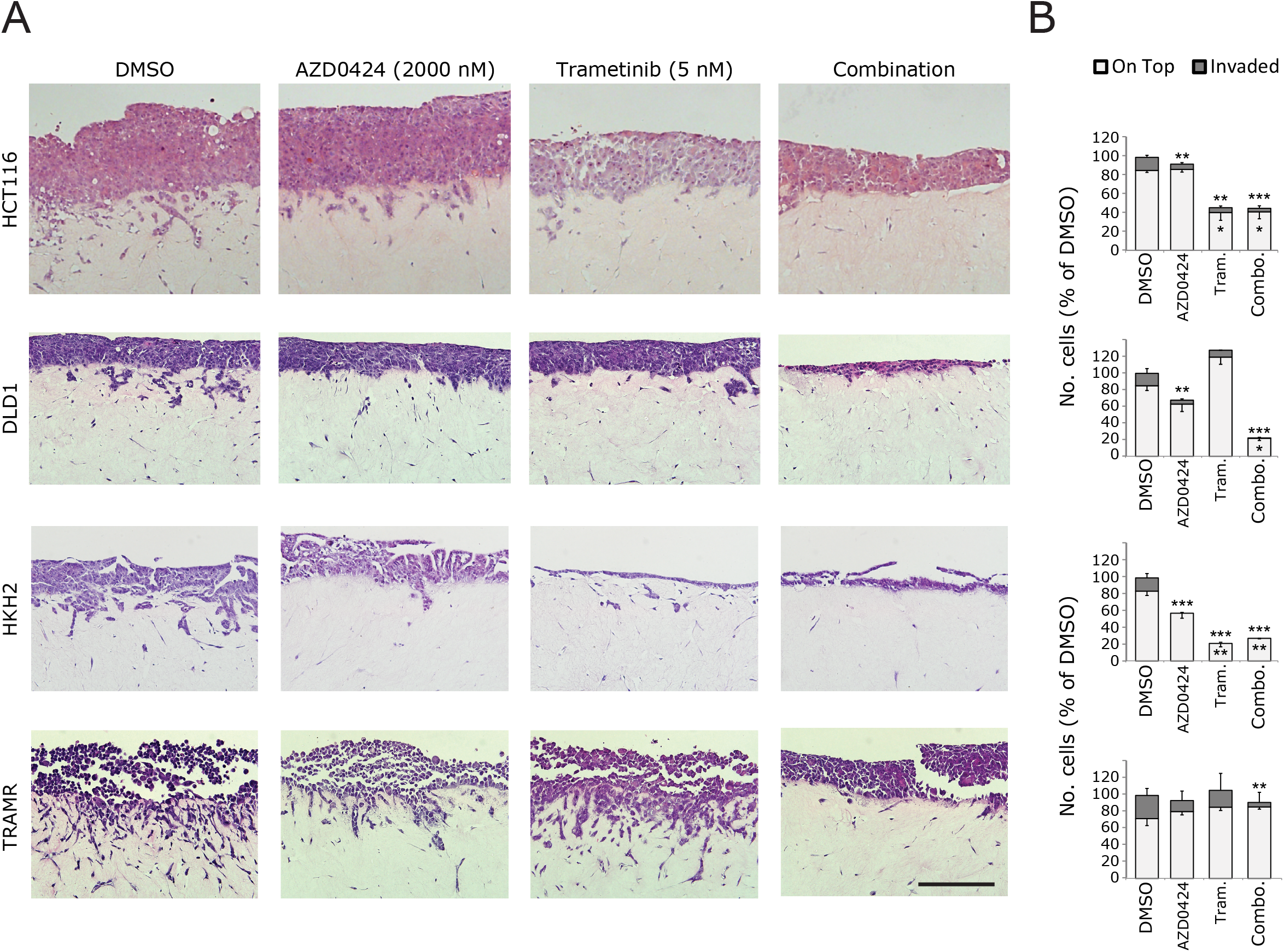
SRC and MEK inhibitor combinations combine to block tumour cell invasion. (A), Organotypic invasion assay. Cells were cultured on top of fibroblast-collagen matrices and allowed to proliferate and invade over a 7-day period with DMSO, AZD0424 (2000 nM), Trametinib (5 nM) or a combination of the two. Images show typical fields of view from H&E-stained sections. (B) Quantification of organotypic invasion. Data are normalised to DMSO values and displayed as means ± SEM (*n* = 3 independent experiments). *, *p* < 0.05; **, *p* < 0.01; ***, *p* < 0.001 (one-way ANOVA).

As KRAS mutation is also known to drive cancer cell invasion and metastasis, we next tested the invasive ability of the HCT116 cells that lack KRAS mutation (HKH2) or that are resistant to MEK inhibitor treatment (TRAMR) cells (Figure 6A and B). As observed in the cell viability assay (Figure 5B), trametinib inhibited proliferation of HKH2 cells but not TRAMR cells. HKH2 cells displayed an epithelial-like morphology and some invasive capacity despite lacking a mutant KRAS signalling while TRAMR cells were morphologically more mesenchymal and more highly invasive (Figure 6A and B). Invasion of HKH2 cell invasion was blocked by either AZD0424 or trametinib treatment while TRAMR cell invasion was only partially sensitive to AZD0424 treatment. However, when these were combined there was further inhibition of invasion of TRAMR cells. Therefore, while proliferation in trametinib-induced resistant TRAMR cells relies on enhanced MAPK pathway activity, and not SRC activity, the switch to a more invasive phenotype is sensitive to AZD0424 alone or combined with submaximal anti-proliferative concentrations of trametinib. Therefore, it is likely that this combination would have maximal effect in reducing invasion of colorectal cancer cells, rather than reducing their proliferation.

## Discussion

Here we evaluated the SRC/ABL kinase inhibitor, AZD0424, and its potential use as an anti-cancer combination therapy by testing across a diverse range of *in vitro* and *in vivo* cancer models in parallel with quantitative pathway profiling at the post-translational level. We sought to characterise how AZD0424, and by extension SRC inhibitors in general, may be employed as part of anti-cancer combination therapy in CRC as AZD0424 treatment could repress transducers of downstream signalling from EGFR, a key driver of metastatic CRC (25, 26). Previous studies indicate that inhibition of MEK in mutant KRAS breast or colorectal cell lines (re)activates many RTKs, sensitising them to RTK targeted therapy (43, 44). *De novo* KRAS mutations reduce the sensitivity of colorectal cells to EGFR targeted therapy (29) and CRC tumours also develop resistance to anti-EGFR therapy by acquiring mutations in RAS (14, 25, 26).

Currently, patients with colorectal cancer are not recommended to receive anti-EGFR therapy if they have mutations in RAS (KRAS, HRAS or NRAS), or BRAF, the exception being when given in combination with drugs (vemurafenib) targeting the BRAF-V600E mutation, in combination with irinotecan and cetuximab (45). Retrospective analyses of clinical trial data has identified that not all KRAS mutations are equal in CRC; KRAS-G13D mutations are sensitive to anti-EGFR therapy (46). Mechanistically, RAS-G13D binds poorly to the RAS-GAP protein NF1 and in cells with hemizygous RAS-G13D mutations (i.e. KRAS-WT/KRAS-G13D), this results in EGFR-dependent activation of RAS-WT; RAS-G12 mutations in contrast bind to and block the activity of NF1 making RAS activation insensitive to anti-EGFR therapy (47). We found that DLD1 cells strongly activated EGFR signalling following MEK inhibitor treatment and that combined trametinib and AZD0424 treatment inhibited cell viability synergistically *in vitro*, but it was not sufficient to block DLD1 tumour growth *in vivo* implying that SRC signalling is dispensible for tumour growth in this model. Indeed, we observed synergistic combination activity upon treatment of DLD1 cells with trametinib and inhibitors of AKT or EGFR family kinases and found that the inhibition of cell viability produced by these combinations was further reduced by the addition of AZD0424 (Supplementary Figure 3). In contrast to DLD1 cells, HCT116 cells do not express NF1 (48) making HCT116 cells unable to activate the EGFR pathway in response to MEK inhibitor treatment. The activation of SRC following MEK pathway inhibition was strongest in HCT116 cells *in vitro* and this was also observed *in vivo* correlating with enhanced inhibition of HCT116 tumour growth upon trametinib and AZD0424 combination treatment relative to respective single agent treatment *in vivo*. Further investigation will be required to test whether CRC cells with RAS-G13D mutations and lacking NF1 represent a subtype sensitive to SRC-MEK inhibitor combinations.

Combinations of SRC and MEK inhibitors have shown benefit in preclinical studies across several tumour types including ovarian, melanoma, non-small cell lung carcinoma, breast and other solid tumours (49-54). In high-grade serous ovarian cancer (HGSOC), combination of MEK (AZD6244) and SRC (saracatinib) inhibitors overcomes EGFR-mediated bypass of the RAS MAPK pathway and targets tumour initiating stem cells (54). Ovarian cancer cells resistant to saracatinib display activation of the MAPK pathway via reduced NF1 expression or overexpression of HER2 and the insulin receptor (52). Mutant-KRAS cell lines are also sensitive to the combination of SRC (dasatinib) and MEK (trametinib) inhibitor treatment by downregulating the Hippo pathway effector TAZ, however, 4 out of 11 cell lines tested were insensitive (53) and so further investigation is required to fully understand the mechanism of this combination and context of cell type sensitivity.

The activation of FAK is a multistep process where first FAK is recruited to the plasma membrane at sites of adhesion by binding PIP2 (55), which primes FAK for autophosphorylation on Tyr397. SRC binds to FAK on Tyr397 and can subsequently phosphorylate other sites on FAK such as the activation loop (Tyr576/577) and other tyrosine residues (Tyr861 or Tyr925) (56). Trametinib treatment of HCT116 cells increased the phosphorylation of FAK on both its autophosphorylation (Tyr397) and SRC phosphorylation (Tyr861) sites (Figure 5). Thus the phosphorylation of FAK on Tyr861 may serve as a biomarker for combined MEK and SRC (or FAK) inhibitors in clinical trials.

Multitargeted inhibitors that reinforce pathway blockade such as VS-6766 which targets both RAF and MEK in the RAS-MAPK cascade can achieve tighter inhibition and reduce pathway reactivation (57). This is an excellent example of the development of a ‘two-drug’ combination in a single compound and this may be an effective strategy for targeting signalling networks supported by SRC. For example preliminary results show that trametinib combined with TPX-0005 (a multitarget kinase inhibitor whose targets include SRC and FAK) synergistically inhibits RAS mutant cell growth *in vitro* and *in vivo* (58). Therefore, Dual EGFR-SRC or FAK-SRC inhibitors could be a future avenue of drug development to address multiple redundant and compensatory signalling mechanisms.

SRC inhibitors have long been recognised as potential anti-invasive/metastatic agents to help improve progression free survival and metastasis free progression(13, 49, 50, 59), yet most clinical trial studies incorporating SRC inhibitors monitor primary tumour growth or regression as a clinical endpoint. Here we have shown that despite either inherent or drug-induced resistance to MEK inhibitors, inhibition of SRC using AZD0424 can effectively block cancer cell invasion in vitro. MEK inhibitor resistance in HCT116 TRAMR cells promoted an aggressive-invasive cell type, most likely driven by their elevated RAS-MAPK signalling and EMT (Figure 6). Biomarkers or pathway signatures of SRC activation following MEK inhibition may predict those patients who would benefit from a SRC-MEK inhibitor combination to combat this aggressive invasive phenotype induced by acquired resistance to MEK inhibitors. Clinical trials in metastatic CRC patients using dasatinib combined with chemotherapy, with or without cetuximab, failed to fully inhibit SRC activity (9). Therefore, efficiently inhibiting SRC activity in a sustainable manner is still a major clinical challenge for the current crop of SRC inhibitors. This could be overcome by the development of novel, well tolerated, highly selective SRC inhibitors (for example (16)). Our data suggest that SRC inhibitors may optimally be combined with other agents to inhibit aggressive invasion in contexts where that is relevant. Clincial trials with appropriate endpoints for metastatic disease would need to be defined.

In conclusion, dynamic signalling networks and pathway switching permit rapid tumour evolution and therapeutic evasion; this requires new and more comprehensive approaches to understand cancer cell signalling networks, ‘driver’ pathways and how best to collapse the robustness of such networks so that tumour cells die in the metastatic niche. Overcoming such dynamic signalling responses may help address high clinical attrition rates associated with target-based drug discovery and improve long term patient outcomes and cancer mortality rates in advanced tumour settings. Complementing advances in Next Generation Sequencing, we have applied protein-level analyses via RPPA to characterise the novel SRC inhibitor AZD0424, including at the post-translational level across a number of cell lines to reveal potential molecular mechanistic insight into compensatory and cooperative mechanisms as well as acquired resistance. We have demonstrated that SRC inhibitors can synergise with MEK inhibitors in colorectal cancer cell lines that depend on RAS-MAPK signalling for survival and invasion, and inhibiting SRC may form part of wider combination regimens that will be most effective when tailored to the pathway activation status of specific patient tumours, and/or to mitigate against enhanced invasion caused by particular therapies such as those targeting MEK.

## Acknowledgements

This work was supported by a Cancer Research UK grant (C157/A1362) awarded to Val Brunton, Neil Carragher and Margaret Frame and by Cancer Research UK Programme Awards (C157/A15703 and C157/A24837) awarded to Margaret Frame and Val Brunton.

