## Supplementary Figure 1 for "Pathway profiling of a novel SRC inhibitor, AZD0424, in combination with MEK inhibitors"

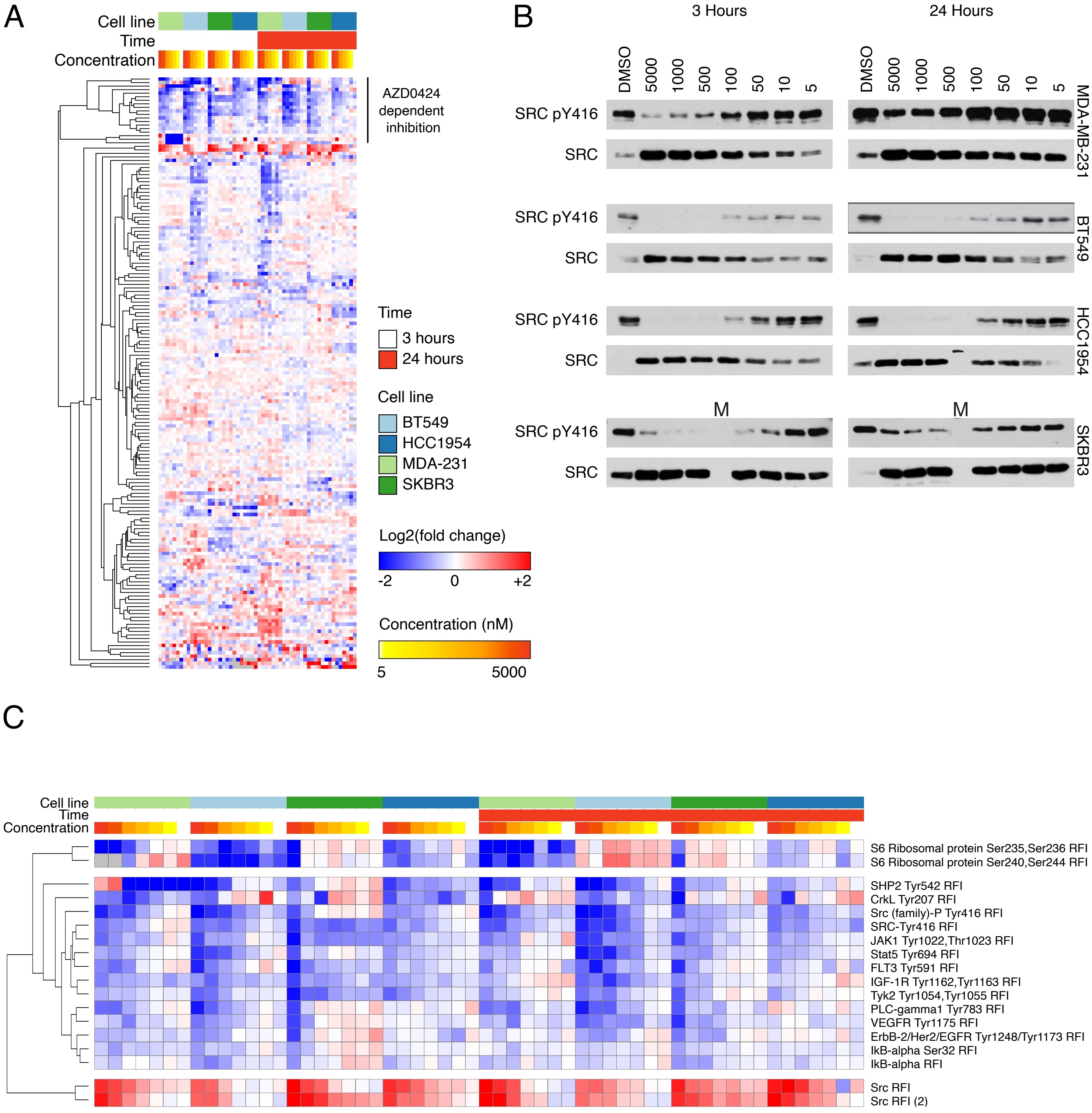


Supplementary Figure 1. RPPA profiling of AZD0424.(A), RPPA profiling of AZD0424 treated breast cancer cell lines. (B), Western blot analysis of cell lysates from breast cancer cell lines treated with AZD0424 probed with anti-SRC and -SRC pY416 antibodies. (C) Cluster of antibodies that display AZD0424 concentraton dependent inhibition.
