## Supplementary Figure 2 for "Pathway profiling of a novel SRC inhibitor, AZD0424, in combination with MEK inhibitors"

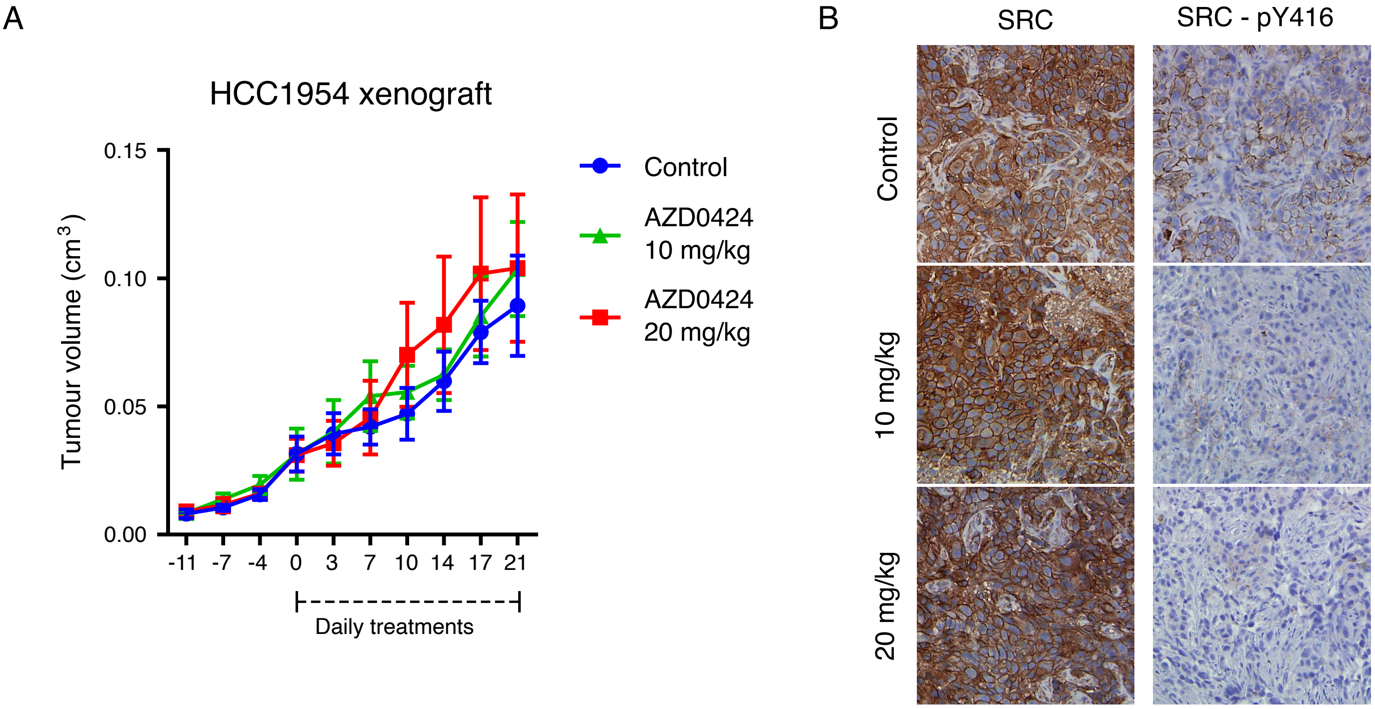


Supplementary Figure 2. AZD0424 inhibits the activation of SRC in vivo. (A), AZD0424 does not inhibit tumour growth of HCC1954 tumours. Tumour volumes are plotted as means ± SEM [n ≥ 4 mice per group (2 tumours per mouse)]. (B), Immunohistochemical analysis of phosphorylated SRC Tyr416 in HCC1954 tumours taken from mice treated with AZD0424.
