## Supplementary Figure 3 for "Pathway profiling of a novel SRC inhibitor, AZD0424, in combination with MEK inhibitors"

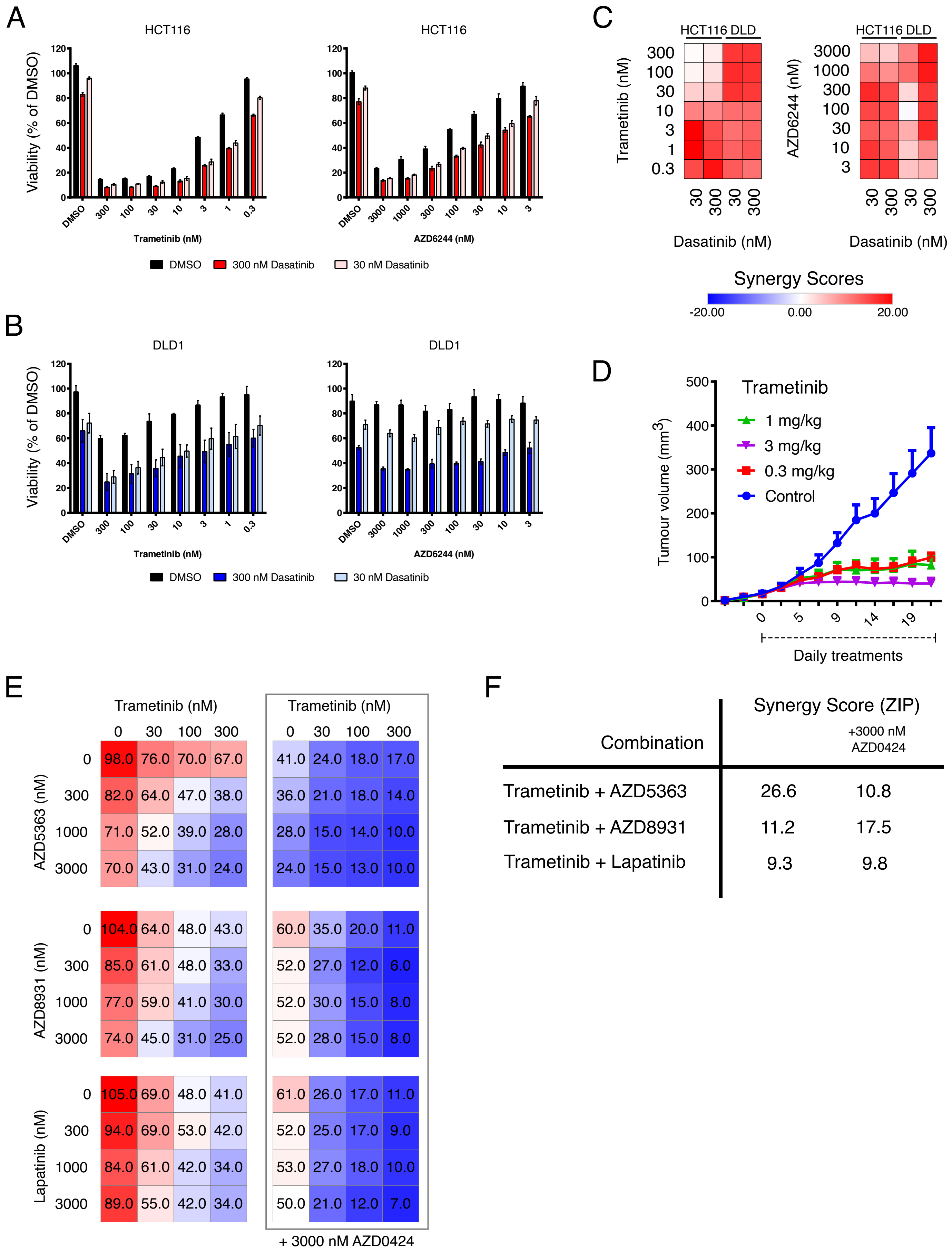


Supplementary Figure 3. SRC and MEK inhibitors synergistically inhibit colorectal cell viability.Dasatinib synergistically inhibits cell viability of HCT116 (A) and DLD1 (B) cells in combination with trametinib or AZD6244. Mean cell viability is shown ± SEM (n = 3 independent experiments). (C) Synergy analysis for trametinib or AZD6244 in combination with dasatinib. Synergy was calculated using the ZIP synergy model. (D) Trametinib inhibits HCT116 tumour growth. Tumour volumes are plotted as means ± SEM [n ≥ 4 mice per group (2 tumours per mouse)]. (E) Inhibitors of AKT (AZD5363) or EGFR family kinases (AZD8931 or lapatinib) synergistically inhibits cell viability of DLD1 cells which can be further enhanced by the addition of AZD0424 as a triple combination. Mean cell viability is shown ± SEM (n = 3 independent experiments). (F) Synergy scores for double and triple combinations.
