## Supplementary Figure 4 for "Pathway profiling of a novel SRC inhibitor, AZD0424, in combination with MEK inhibitors"

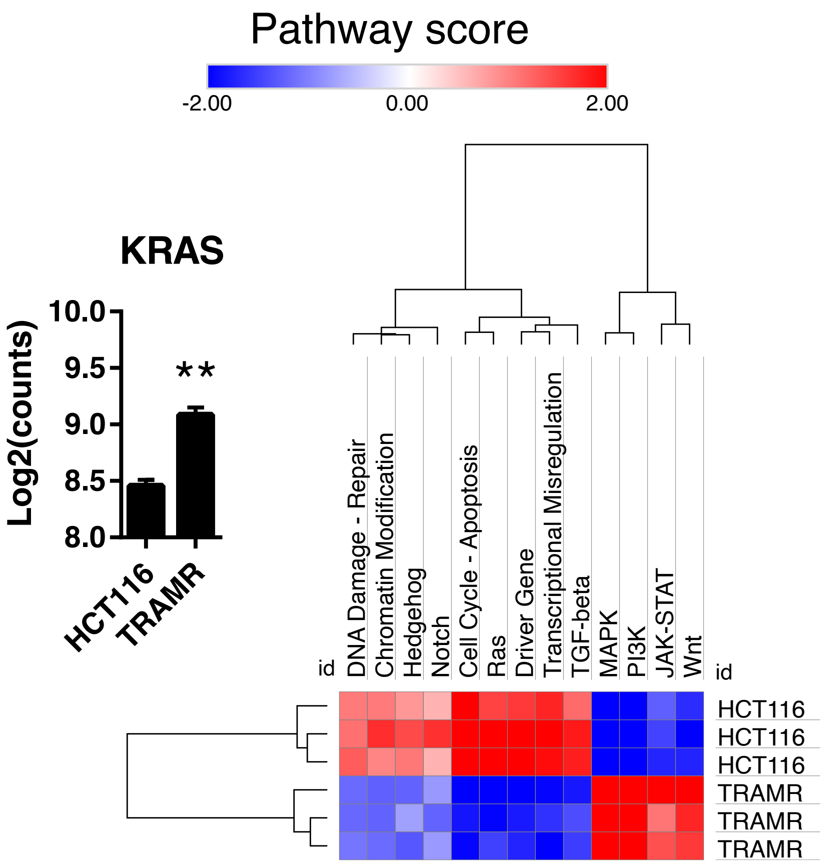


Supplementary Figure 4. MAPK signalling is overactivated in trametinib resistant cells.RNA from HCT116 and trametinib resistant TRAMR cells was analysed using a NanoString PanCancer Pathway panel. Inset bar chart shows the expression of KRAS gene. Mean cell expression is shown ± SEM (n = 3 independent experiments). **, $p < 0.01$ (t-test).
