## Supplementary Figure 5 for "Pathway profiling of a novel SRC inhibitor, AZD0424, in combination with MEK inhibitors"

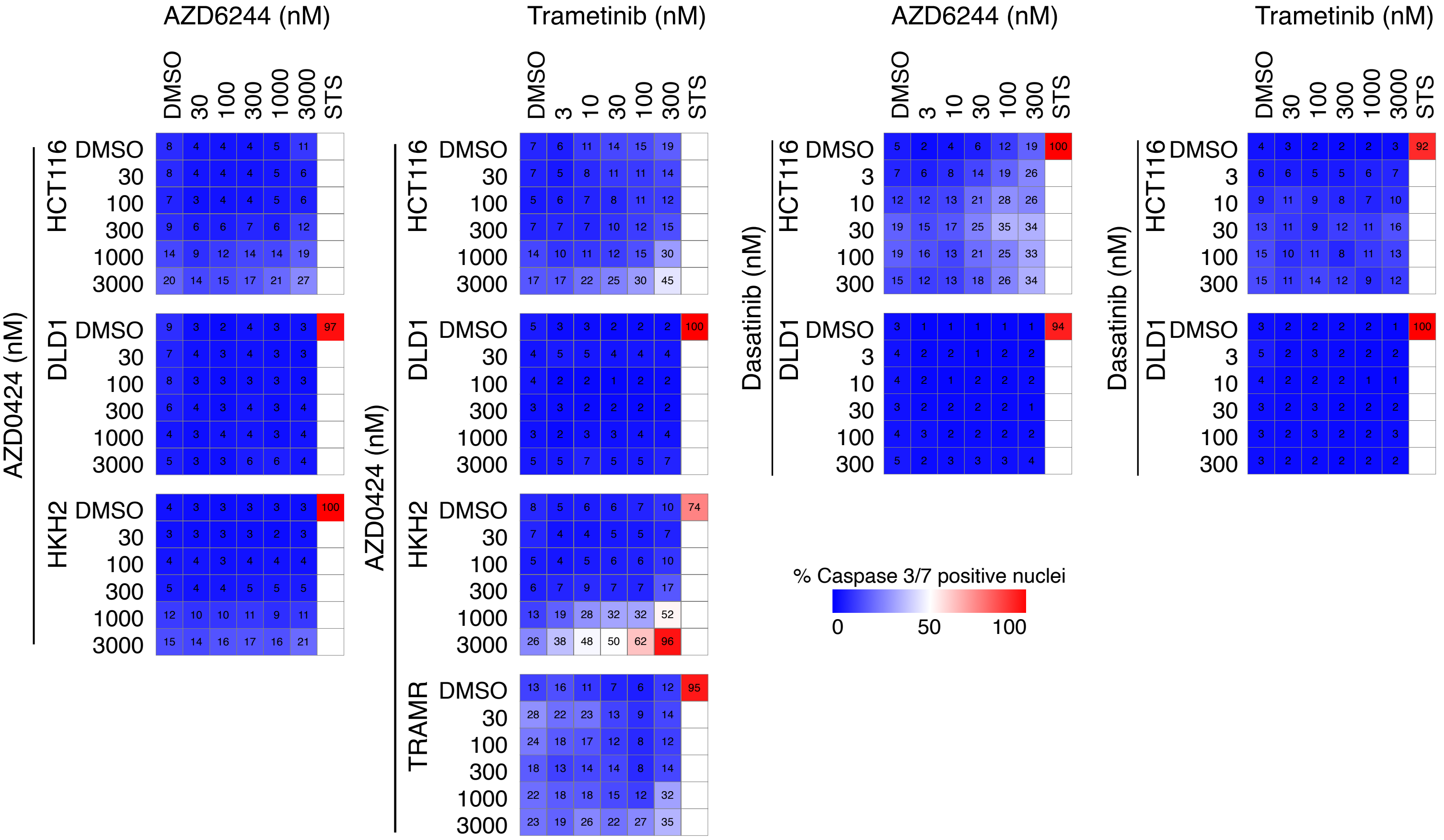


Supplementary Figure 5. The combination of SRC and MEK inhibitors only weakly induces apoptosis in colorectal cell lines.

Activation of caspase 3/7 in cells (HCT116, DLD1, HKH2 or TRAMR) following 48 hours treatment with SRC inhibitors (AZD0424 or dasatinib) with MEK inhibitors (AZD6244 or trametinib) in combination.
